# Tuberculosis treatment failure associated with evolution of antibiotic resilience

**DOI:** 10.1101/2022.03.29.486233

**Authors:** Qingyun Liu, Junhao Zhu, Charles L. Dulberger, Sydney Stanley, Sean Wilson, Eun Seon Chung, Xin Wang, Peter Culviner, Yue J. Liu, Nathan D. Hicks, Gregory H. Babunovic, Samantha R. Giffen, Bree B. Aldridge, Ethan C. Garner, Eric J. Rubin, Michael C. Chao, Sarah M. Fortune

## Abstract

Antibiotics are a cornerstone of medicine, placing bacterial pathogens under intense pressure to evolve new survival mechanisms. Analysis of 51,229 *Mycobacterium tuberculosis (Mtb)* clinical isolates identified an essential transcriptional regulator, *Rv1830 (*here named *resR)* as a frequent target of positive (adaptive) selection. *resR* mutants do not demonstrate canonical drug resistance or drug tolerance but instead have significantly faster recovery after drug treatment across all antibiotics and combinations tested, a phenotype which we term antibiotic resilience. ResR acts in a regulatory cascade with other growth-controlling transcriptional regulators WhiB2 and WhiA, which are also under positive selection in *Mtb* clinical isolates. Mutations of these genes are associated with treatment failure and the acquisition of canonical drug resistance.

---

Antibiotics are designed to kill bacteria by inhibiting biological processes essential for survival. Completely eliminating a population of bacteria with antibiotics is challenging even when the bacteria are nominally sensitive to the drug (*1*). There is a growing body of work defining the mechanisms beyond canonical resistance by which bacterial populations avoid clearance by antibiotics (*2, 3*). These include the formation of privileged subpopulations of drug-tolerant bacteria, as well as the acquisition of other traits that enhance bacterial survival, including variation in the cellular requirements for the target and alterations in drug penetration or efflux (*2-4*). We hypothesized that bacterial adaptation to antibiotics would be recorded as mutations in the genes associated with the most clinically relevant of these adaptive processes, and dissecting the targets of natural selection would reveal new mechanisms that enable bacteria to survive antibiotics in patients. Therefore, we sought to identify the evolutionary signatures of adaptation to antibiotics in *Mycobacterium tuberculosis* (*Mtb*), an obligate human pathogen that has remained one of the largest causes of mortality, despite facing months of antibiotic pressure in every treated patient and which, as a global population, has been under antibiotic selection since the introduction of streptomycin in 1944 (*5*).

## Ongoing positive selection in Mtb population

When clinical *Mtb* isolates are sequenced, the specimen is typically not derived from a single colony but rather the bacterial population sampled from the patient. This population contains mutations arising from the within-host evolution of the bacterium, which can be tracked by identifying unfixed mutations via deep sequencing (Fig. 1A) (*6, 7*). We reasoned that by looking at very large numbers of clinical isolates, even in the absence of clinical metadata, we could identify genes that repeatedly acquire mutations (parallel evolution) within different patients which would mark processes under positive selection. Therefore, we assembled previously published whole-genome sequencing data from 51,229 *Mtb* clinical isolates (Fig. 1B) and used a mutation burden test and the ratio of nonsynonymous to synonymous mutations (*dN/dS*) to identify genes that are under positive selection in the *Mtb* population (Materials and Methods). We also identified intergenic regions (IGRs) that are highly enriched for mutations. As expected, genes and IGRs previously associated with drug resistance or tolerance are highly mutated and the genes have *dN/dS* ratios >1, indicative of positive selection (Fig. 1B, fig. S1). Interestingly, we also identified a second set of genes that show similar selective signals, but for which the selective pressures are unknown (Fig. 1B).

**Fig. 1.**
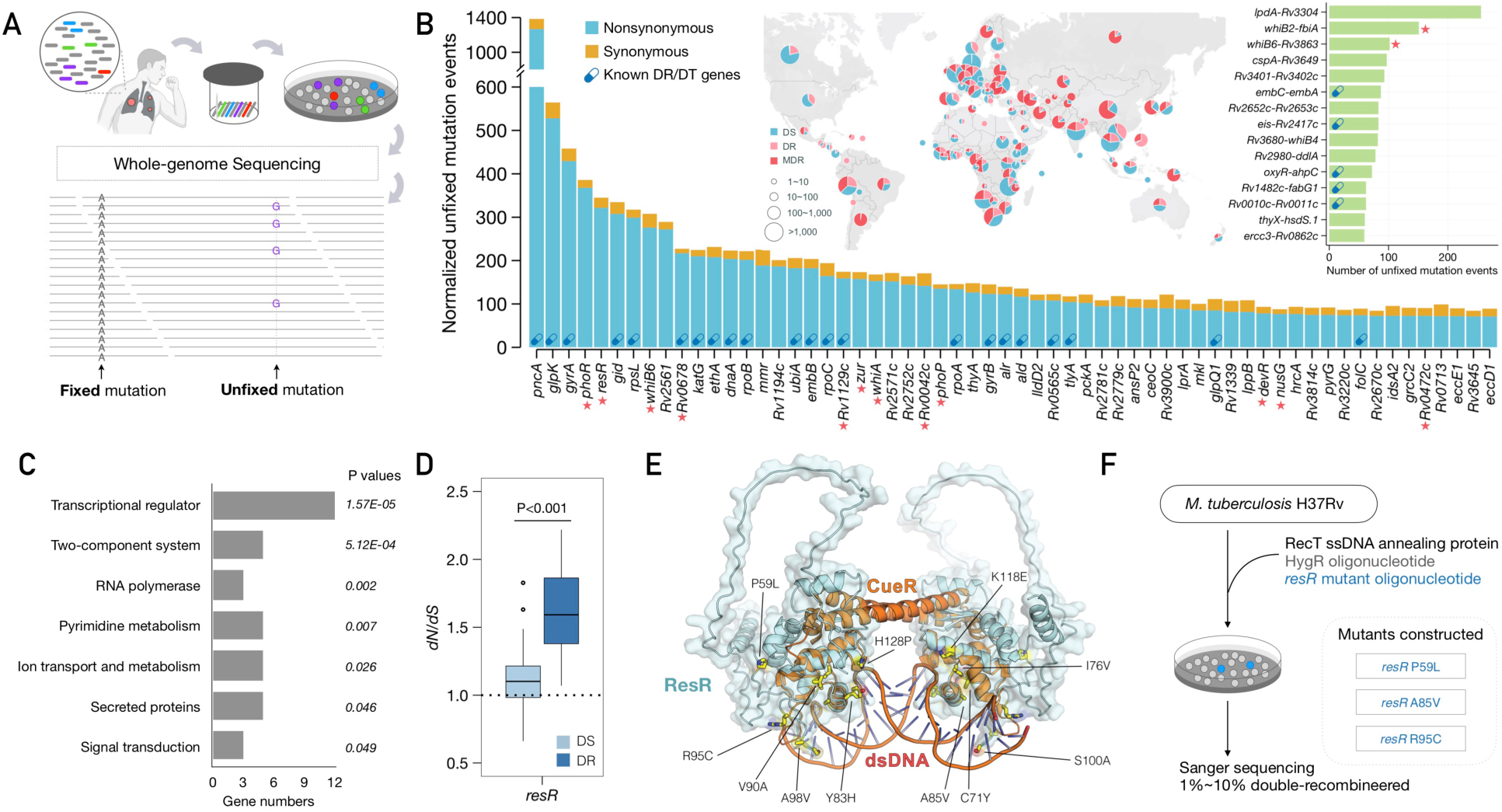
Ongoing positive selection in the global *Mtb* population. (**A**) Diagram of fixed and unfixed variants. (**B**) Genes and IGRs with signal of ongoing positive selection. Known drug-resistance (DR) or drug-tolerance (DT) genes are marked with capsule symbols. Red stars indicate transcriptional regulators. The map shows the geographic origin of the clinical *Mtb* isolates and their DR status; DT (drug-tolerance), MDR (multi-drug resistance). (**C**) Categories of genes under positive selection, with adjusted *P* values for enrichment. (**D**) dN/dS ratio of *resR* in DR and DS strains, *P* value by student *t* test. (**E**) Structural model of the ResR dimer (cyan) aligned to the MerR-family homolog CueR (orange) in complex with duplex DNA with common ResR clinical mutations indicated (yellow). (**F**) Schematic of construction of mutations in the chromosomal copy of *resR* in *Mtb*.

Functional enrichment analysis indicated that the genes under selection are highly enriched for transcriptional regulators (12/60, P=1.57E-5, Fig. 1B, C). One of the most frequently mutated regulators is *resR (*gene identifier: *Rv1830)*, an essential gene that is predicted to be a *merR*-type regulator. We also found evidence of positive selection in an analysis of the fixed mutations in *resR* (*dN/dS* = 2.91); the pattern of mutation is very similar between unfixed and fixed mutations (fig. S2), suggesting that within-host selection leads to mutation fixation. Importantly, we found the *dN/dS* of *resR* is higher in drug-resistant strains as compared to that in drug-sensitive strains (Fig. 1D), suggesting that *resR* mutations are associated with the evolution of drug resistance. Protein structural modeling indicates that most *resR* clinical mutations occur in the *merR*-type helix-turn-helix DNA binding domain with the most common mutations located at the interface between ResR and duplex DNA (Fig. 1E and fig. S2), where they could affect DNA binding affinity and the transcription of downstream genes.

## resR mutants display “antibiotic resilience”

To investigate the functional consequence of *resR* mutations, we used oligo-mediated recombineering to make point-mutant strains by introducing three common clinical mutations (P59L, A85V and R95C respectively) into the chromosomal copy of *resR* in *Mtb* (H37Rv) (Fig. 1F). The *resR* mutations did not affect bacterial growth under standard or host-relevant carbon source conditions (fig. S3). However, we noted that *resR* mutants are about 20% longer and about 5% wider than the wild-type cells (fig. S4), and have a thickened cell envelope (fig. S5), suggesting that mutations of *resR* have functional consequences under standard growth conditions.

We next tested the effects of *resR* mutations on drug resistance by measuring minimum inhibitory concentration (MIC) for a panel of 8 antibiotics including first-line, second-line and new anti-tuberculosis (TB) drugs (Fig. 2A). *resR* mutants and wild-type strains have very similar MIC profiles, with only subtle MIC shifts identified for three drugs -- a small increase for isoniazid (INH, 0.01µg/mL to 0.02µg/mL) and minor decreases for rifampicin (RIF, 0.0028µg/mL to 0.0024µg/mL) and ofloxacin (OFX, 0.11µg/mL to 0.098µg/mL) (Fig. 2A, fig. S6). These MIC changes are far below the MIC for classification as drug-resistant (Fig. 2A). We next assessed the effects of *resR* mutations on drug tolerance by performing time-kill analyses for the 8 antibiotics, using drug concentrations at 100X of the MICs. The *resR* mutants and wildtype strains show very similar time-kill dynamics although again there are subtle differences for some drugs (OFX, ETH and BDQ) at particular time points (Fig. 2B). The changes in the minimum duration of time to kill 99% of bacteria (MDK_99_) are less than 24 hours (Fig. 2B); previously characterized drug-tolerance mutants in *Mtb* prolong the MDK_99_ by more than 4 days (*8*). Thus, we conclude that *resR* mutations do not have a major impact on either canonical drug resistance or drug tolerance.

**Fig. 2.**
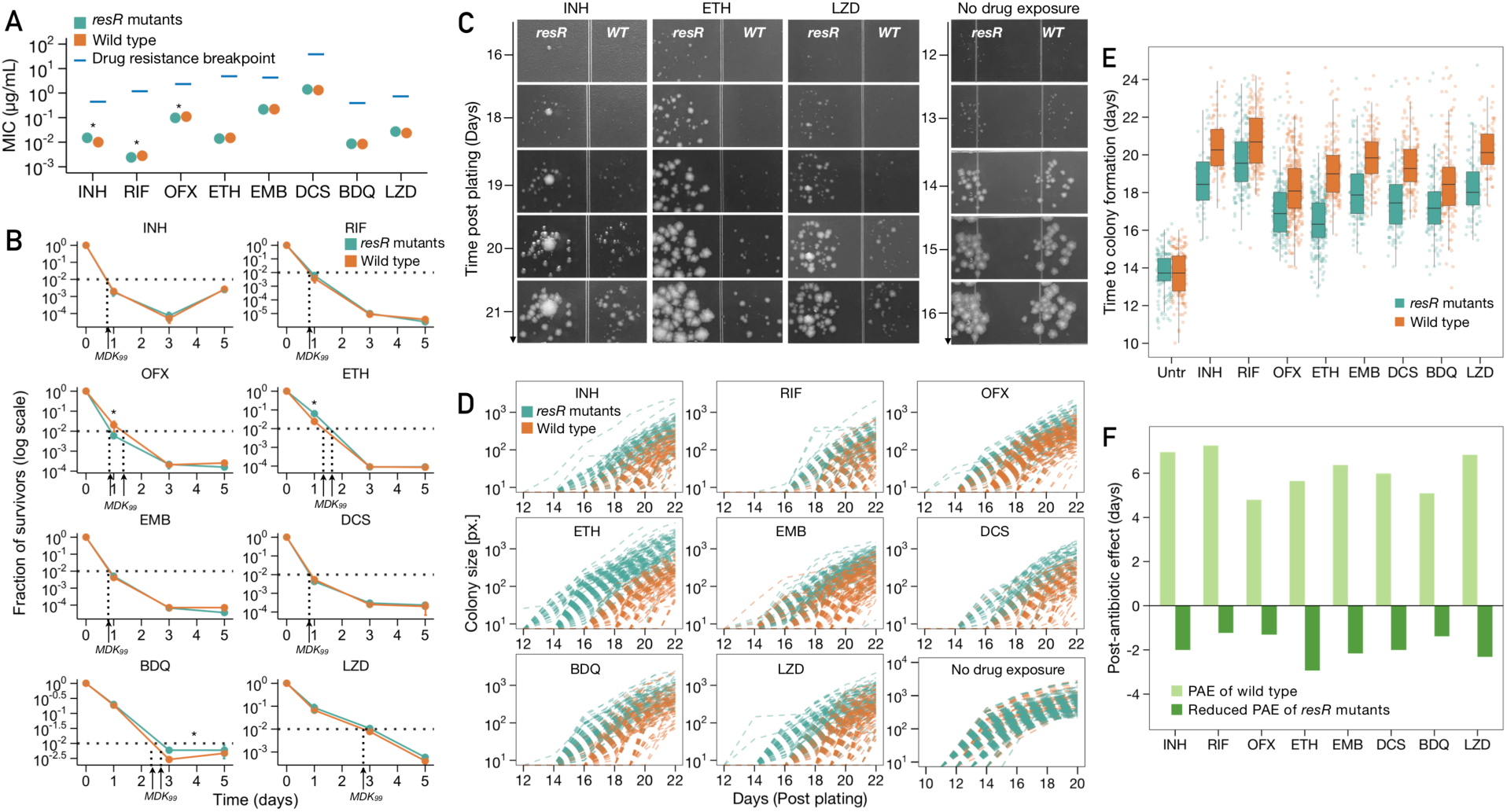
*resR* mutants showed faster recovery after drug exposure. (**A**) MICs of *resR* mutants and wild-type for 8 anti-TB drugs. INH: Isoniazid; RIF: Rifampicin; OFX: Ofloxacin; ETH: Ethionamide; EMB: Ethambutol; DCS: D-cycloserine; BDQ: Bedaquiline; LZD: Linezolid. Drug resistance breakpoint is the drug concentration that defines clinical drug resistance. (**B**) Time-kill kinetics for 8 different drugs of *resR* mutant and wild-type *Mtb* strains. The concentration of each drug was used as 100-fold of MIC. * symbol indiates significance by unparied *t* test. (**C**) Representative images illustrating the post-antibiotic recovery dynamics of *resR* mutant and wild-type (WT) *Mtb* strains. (**D**) Quantitative colony-size tracking in pixel unit (px.) for *resR* mutant and wild-type *Mtb* strains in the presence or absence of antibiotic exposure. (**E**) Duration from plating to the appearance of visible colonies (Time to colony formation) for *resR* mutant and wild-type strains after exposure to indicated antibiotics or no drug exposure (Untr). *P=0*.*7291* for Untr group and *P<0*.*0001* for all drug groups, by Mann-Whitney test. (**F**) The median value of post-antibiotic delay of wild type (light green) and the median value of reduced post-antibiotic delay of *resR* mutants (dark green).

In the time-kill assays, however, we observed an unexpected phenotype that clearly distinguished *resR* mutants from the wild-type strains. After the cells were plated on drug-free media post antibiotic exposure, *resR* mutants formed visible colonies significantly earlier than the wild type (Fig. 2C). This early recovery phenotype of *resR* mutants was observed for all the 8 antibiotics tested while no difference was seen in the absence of drug exposure (Fig. 2C, fig. S7). Using quantitative image analysis to track the growth of individual colonies over time (Fig. 2D, fig. S8) (*9*), we found that antibiotic treatment causes a 4.7∼6.8 day delay in colony formation for wild-type strains as compared with the “no drug exposure” group, which is highly reproducible and characteristic for each drug (Fig. 2E). This delay – classically referred to as the post-antibiotic effect (PAE) - has been recognized for decades across many bacterial pathogens and is an important factor in the design of treatment regimens (*10, 11*). The PAE (indicated as time to colony formation) is reduced in *resR* mutants by 20∼50% for all 8 drugs tested, with an average reduction of 1.9 (1.24∼2.94) days (Fig. 2E, F). The *resR* mutants also showed faster recovery when challenged by combinations of the first-line anti-TB drugs (fig. S9).

To date, the PAE remains mechanistically enigmatic with a recent work favoring a detoxification model in which the cells must export residual drug before growth resumes (*12*). These results suggest that *Mtb* has mechanisms for post-antibiotic recovery that are under genetic control and that *resR* mutants optimize this system to accelerate recovery after a wide variety of antibiotic insults. We term this phenotype antibiotic resilience, and name this gene (Rv1830), *resR* for <u>Res</u>ilience <u>R</u>egulator.

## Antibiotics resilience manifests as “early wake up”

To better understand early events in post-antibiotic recovery, we used microscopy to interrogate cell regrowth after antibiotic wash-off. *Mycobacteria* grow from their subpolar regions, and this outgrowth can be quantitatively defined via imaging by tracking the incorporation of fluorescently labeled D-amino acids (FDAA) into the nascent peptidoglycan near the cell poles (*13, 14*). We assessed bacterial regrowth after 24 hours of INH exposure at 100X MIC, followed by a 10-36 hour recovery window in drug-free media (Fig. 3A, B). During the recovery period, *resR* mutants demonstrate significantly more new cell wall synthesis than the wildtype, as early as 24 hours post recovery (P<0.0001 for 24h and 36h, Fig. 3B, C). We also used flow cytometry to rapidly and quantitatively assess new cell wall synthesis in mutant and wild-type cells after exposure to the other antibiotics, and again found that *resR* mutants show increased cell wall synthesis during regrowth after exposure to all 8 antibiotics tested (Fig. 3D) and the different *resR* mutants showed similar phenotype (fig. S9D). Wild-type cells and *resR* mutants have similar FDAA incorporation in the absence of antibiotic exposure (Fig. 3D).

**Fig. 3.**
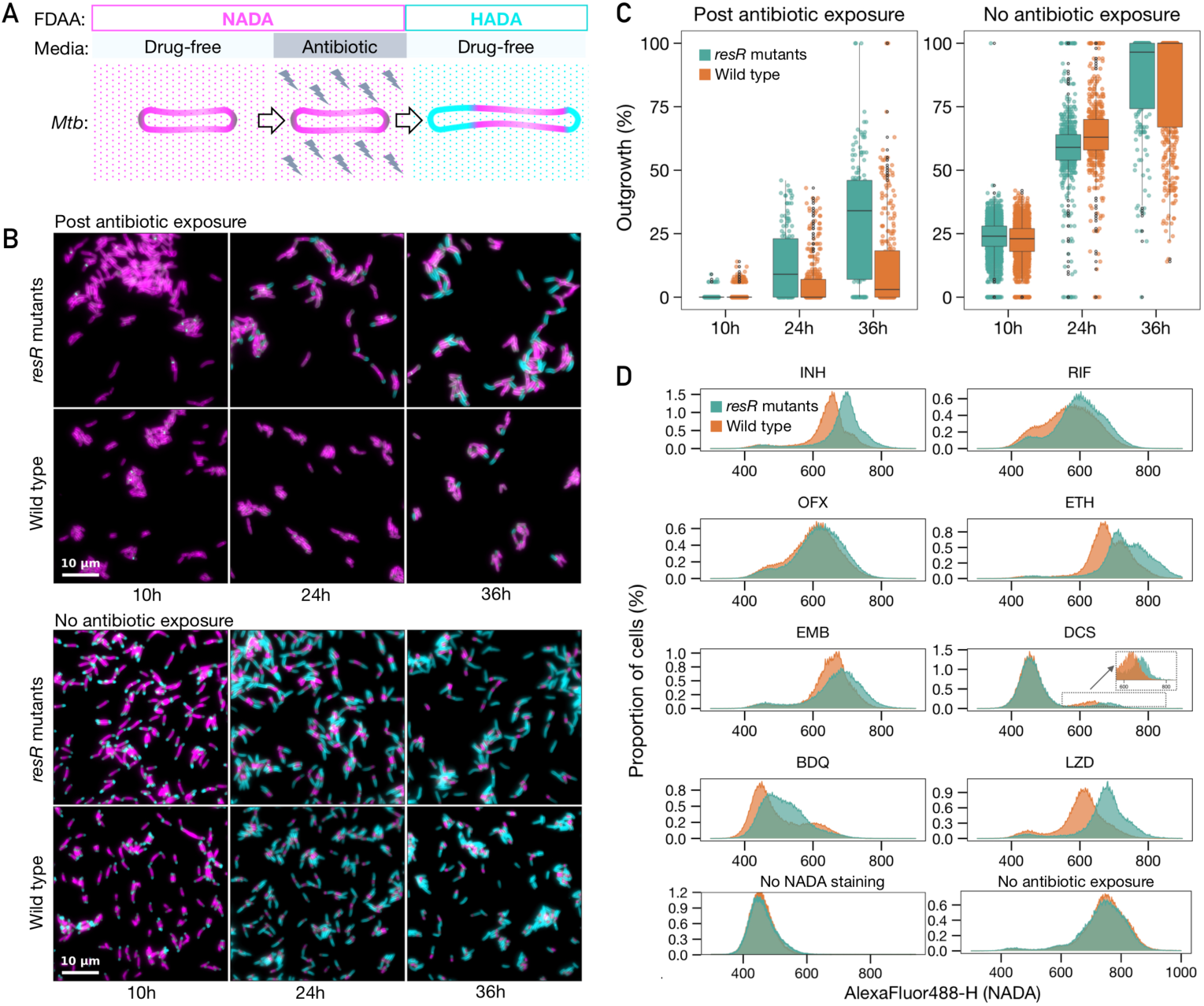
Antibiotic resilience manifests as early regrowth. (**A**) A schematic diagram of pulse-chase experiment for tracking the regrowth of *Mtb* cells post antibiotic exposure as tracked by fluorescent d-amino acid incorporation (NADA and HADA) (**B**) Microscopy of the *resR* mutants and wild type during regrowth after 24 hours of INH (100X MIC) exposure (the upper panel) and no antibiotic exposure (the lower panel). (**C**) Quantitative comparison of outgrowth between *resR* mutant and wild type in post antibiotic exposure group and no antibiotic exposure group, *P*<0.0001 for 24h and 36h of Post antibiotic exposure group by Mann Whitney test. (**D**) Flow cytometry for NADA incorporation into *Mtb* cells after 24 hours recovery post antibiotic exposure or without drug exposure.

## A regulatory pathway underlying antibiotic resilience

To delineate the regulatory targets through which ResR mediates antibiotic resilience, we first used *in vitro* whole-genome DNA binding and deep sequencing assay (IDAP-Seq)(*15*) to identify the ResR binding sites across the genome. We purified ResR proteins with either an N-terminal His tag or C-terminal His tag, and performed IDAP-Seq with both versions. We identified four major ResR binding sites in the *Mtb* genome: the intergenic regions between *rimJ-Rv0996, nrdH-Rv3054, whiB2-fbiA, and Rv3916c-parB* (Fig. 4A).

**Fig. 4.**
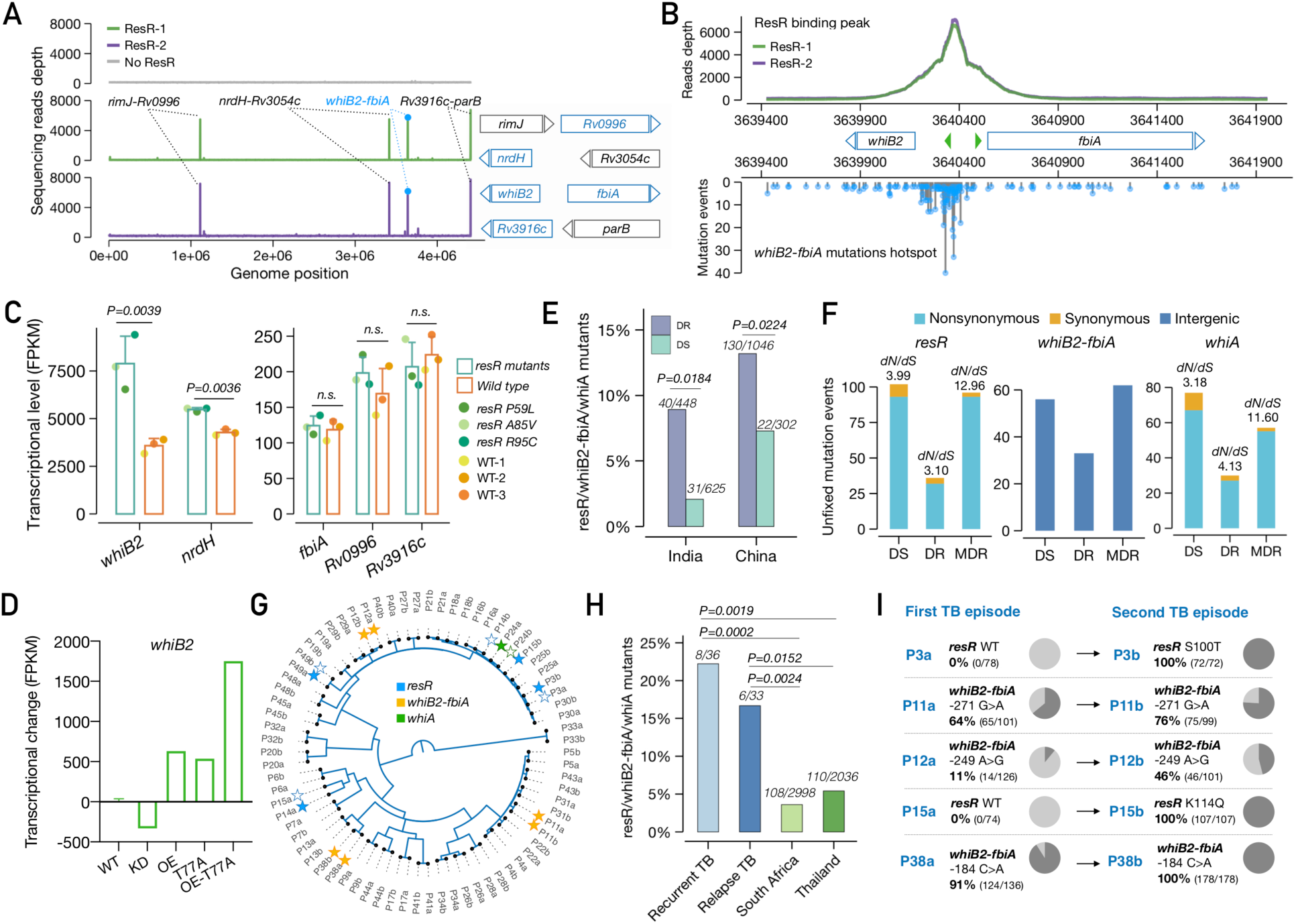
ResR activates *whiB2* and mutations in the pathway were associated with relapse of drug-susceptible tuberculosis. (**A**) IDAP-Seq identified binding sites of ResR. (**B**) ResR binding peak overlaid with the mutations identified in clinical strains between *whiB2-fbiA*. The transcriptional start sites of *whiB2* and *fbiA* are annotated (green arrows). (**C**) Expression of putative ResR targets in *resR* mutants and wild-type strains, *P* values by paired *t* test. FPKM: fragments per kilobase of exon per million mapped fragments. (**D**) Transcriptional changes of *whiB2* in *M. smegmatis* strains: CRISPR-i knock down (KD); over expression (OE); point mutant (T77A); over-expression of point mutant (OE-T77A). (**E**) The proportion of *resR, whiB2-fbiA* and *whiA* mutants in DR and DS strains sequenced from India and China, *P* values by Fisher’s exact test. (**F**) Unfixed mutations in DS, DR and MDR (resistant to RIF and INH) strains. (**G**) A phylogenetic tree of paired *Mtb* isolates from 36 recurrent TB patients. Solid stars indicate isolates in which mutations were detected while empty stars indicate absence of mutations in one of the paired isolates. (**H**) Percentage of isolates with mutations in *resR/whiB2-fbiA/whiA*. “Recurrent TB” includes 3 patients with *Mtb* isolates suggestive of re-infection. (**G**) Mutational trajectory in *Mtb* isolates from the first TB to second TB episodes in 5 pairs of isolates. The mutations in other three pairs: P14 (*resR*, D144N; 100% in P14a, 0% in P14b); P49 (*resR*, R95H; 8.7% in P49a, 0% in P49b); P24 (*whiA*, A131T; 11.6% in P24a, 0% in P24b)

None of the genes flanking the dominant ResR binding sites have a known role in drug resistance. Surprisingly, however, our population genomic analyses found that the *whiB2-fbiA* intergenic region is also one of the major targets of ongoing positive selection in clinical *Mtb* isolates (Fig. 1B). Moreover, the site where ResR binds in the *whiB2-fbiA* intergenic region is precisely the same site where mutations are accumulating in clinical *Mtb* strains (Fig. 4B), an evolutionary pattern suggesting natural selection has acted both the regulator protein and its binding site, presumably to alter the expression of *whiB2* or *fbiA (*Fig. 4A).

To further investigate the regulatory role of ResR for these potential targets, we used RNA-Seq to assess gene expression changes in three independent *resR* mutants. The *resR* mutants consistently show increased *whiB2* transcription and a modest increase in *nrdH* expression but no significant alterations in the expression of the other genes adjacent to the major ResR binding sites (Fig. 4C). The changes in *whiB2* expression were part of a pattern of altered transcription that also included increased expression of the *iniBAC* operon, previously shown to be expressed in response to cell wall targeting antibiotics (*16*); several PPE genes including *PPE51* implicated in trehalose transport(*17*); and other mediators/regulators of central carbon metabolism (*bkdABC, icl1, clgR*) and cell wall acting genes, notably *pbpB(18, 19)*.

To further define the regulatory relationship between *resR* and *whiB2*, we reconstructed several *resR* variants in *Mycobacterium smegmatis* (homolog: *MSMEG_3644*), including CRISPR-i knockdown, overexpression and also point-mutant strains (Fig. 4D). RNA-Seq of these strains suggests that ResR functions as an activator of *whiB2* expression and that the clinical mutation (T77A) result in increased *whiB2* transcription (Fig. 4D). WhiB2 is an essential regulator that controls cell growth division in mycobacteria; knockdown of *whiB2* expression results in cell elongation and division defects, while overexpression of *whiB2* in *Mtb* results in increased cell length (*20, 21*), which is consistent with the phenotype of *resR* mutants (fig. S4). The regulon of WhiB2 has been characterized in other actinobacteria like *Streptomyces* where the WhiB2 homolog interacts with another regulator, WhiA, to initiate cell division (*22, 23*). Strikingly, *whiA*, like *resR* and *whiB2*, is also a major target of positive selection in clinical *Mtb* isolates (Fig. 1B).

## resR, whiB2-fbiA and whiA mutants are associated with acquisition of canonical drug resistance and TB relapse

To assess the clinical relevance of mutations in this regulatory triad, we compared the prevalence of fixed mutations of *resR, whiB2-fbiA* and *whiA* in drug-susceptible (DS) strains versus drug-resistant (DR) strains. Mutations in these genes/IGR are significantly more frequent in DR strains as compared to DS strains in data from all countries (fig. S10A). We further looked at *Mtb* strains from high TB burden countries, India and China, where *Mtb* populations were not skewed by recent expansions of clonal *Mtb* strains and each with >1,000 sequenced isolates. Again, strains with the three genes/IGR’s mutations have increased odds ratios for drug resistance (India, OR: 1.80; 95% CI: 1.11∼2.91; China, OR: 1.71; 95% CI: 1.06∼2.74) (Fig. 4E). Therefore, these data suggest that *resR, whiB2-fbiA* and *whiA* mutations facilitate the evolution of canonical drug resistance. Importantly, we noted that *resR, whiB2-fbiA* and *whiA* remain under positive selection in DR strains and even multidrug-resistant (MDR) strains, as assessed by within-host evolution (Fig. 4F), suggesting that even after the emergence of resistance to first-line drugs, these mutations continue to provide an advantage for the bacterium as *Mtb* encounters second-line agents.

Finally, while *resR, whiB2-fbiA* and *whiA* variants are more frequent in DR strains, variants in these genes are also under positive selection in DS strains (fig. S10B) and the sites of mutation in DS strains mimic those in DR strains (fig. S10C). Indeed, 1.5%-9.7% of DS *Mtb* strains from the high TB burden countries have fixed mutations in one of these three genes/IGR (fig. S10D). We postulated that in addition to being clinically important as stepping-stone mutations for the emergence of canonical drug resistance, these variants independently contribute to treatment failure in DS patients. To test this hypothesis, we reanalyzed data from the REMoxTB trial, a global Phase 3 clinical trial that sought to reduce the treatment duration of DS TB (*24, 25*). Of the 1931 patients enrolled in the trial, paired *Mtb* isolates from 36 recurrent TB patients were whole-genome sequenced and the isolates remained drug-sensitive (*25*). We found 8/36 (22.2%) of the isolates from patients who failed treatment had *resR, whiB2-fbiA or whiA* mutations (Fig. 4F, fig.S10E), a frequency significantly higher than found in the background populations of South Africa (108/2998, 3.6%) and Thailand (110/2036, 5.4%) where the trial was conducted (P=0.0024 and P=0.0152 respectively, Fisher’s exact test, Fig. 4G). Additionally, the within-host frequency of these variants increased between the initial and recurrent episodes, in three cases rising to fixation (Fig. 4H).

Taken together, we have identified a new form of altered drug susceptibility, antibiotic resilience, which is regulated by *resR;* mutations in the regulating cascade are positively selected in clinical *Mtb* isolates and associated with poor treatment outcomes in TB. Current clinical measures of antibiotic susceptibility focus largely on the magnitude of drug exposure – what concentration of drug the pathogen can experience and grow or at least survive (*26*). However, our results suggest that in the clinic, *Mtb* is evolving to change the temporal dynamics of its recovery after drug exposure. The temporal dynamics of drug responses are considered when developing new drugs and drug regimens but often forgotten as potential drivers of treatment outcomes that might be influenced by pathogen variation. Interestingly, in studies of *E. coli* experimentally evolved under intermittent drug exposure, temporal variants were the first to emerge (*27*). In this study, the mutants had delayed regrowth rather than faster regrowth, perhaps because the temporal phenotypes that emerge vary by the pathogen, drug regimen and host factors such as drug metabolism (*28-30*). As current clinical diagnostics are largely blind to temporal phenotypes, selection for temporal phenotypes such as antibiotic resilience may underlie treatment failure in other drug-susceptible infections.

## ACKNOWLEDGMENTS

We thank Timothy M. Walker for collecting and sharing the information of the geographic origin of more than 10,000 *Mtb* isolates. We also thank Christopher M. Sassetti for the helpful discussions during this work.

## Author contributions

Q.L., J.Z., C.L.D., S.S., S.W., C.C., X.W., P.C., J.L., N.D.H., G.H.B. and S.R.G. performed experiments and data analysis. B.B.A., E.C.G., E.J.R., M.C., S.M.F supervised this study. Q.L., J.Z., M.C. and S.M.F wrote the manuscript.

## Data and materials availability

All data are available in the main text or the supplementary materials and sequencing data are deposited in the Sequence Read Archive database (PRJNA820190).

