## Supplementary figures for "Tuberculosis treatment failure associated with evolution of antibiotic resilience"

**Fig.S1 - Fig.S10**

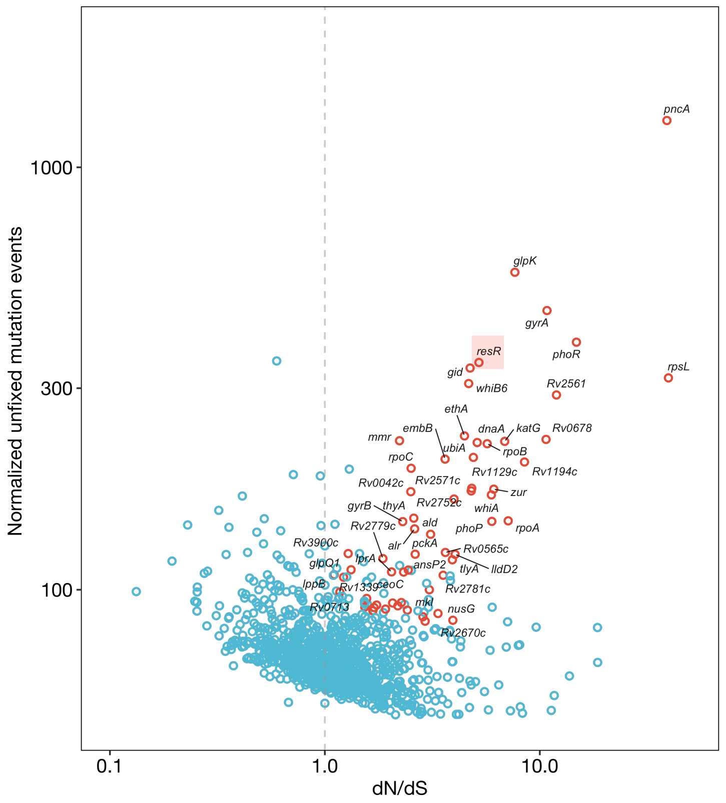


**Fig. S1. Gene candidates under ongoing positive selection in *Mtb* population.** A bubble plot showing the normalized unfixed mutation events and *dN/dS* ratio of *Mtb* genes, with the top 60 gene candidates shown in red and the remaining in blue. *resR* is highlighted in red background.

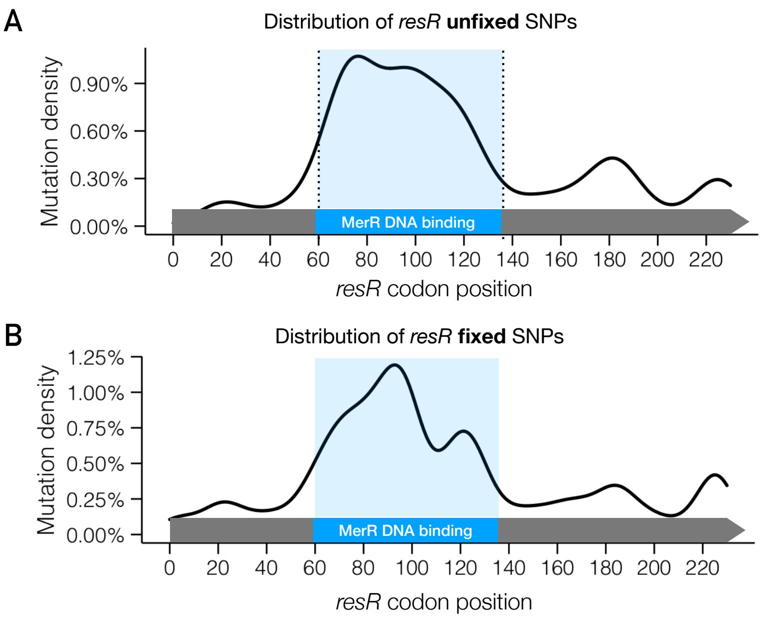


**Fig. S2. Mutation hotspots of unfixed and fixed SNPs of *resR*.** Mutation distribution of unfixed SNPs (**A**) and fixed SNPs (**B**) in gene *resR*. The MerR-type DNA binding domain (codon 60 – codon 136) is highlighted in blue.

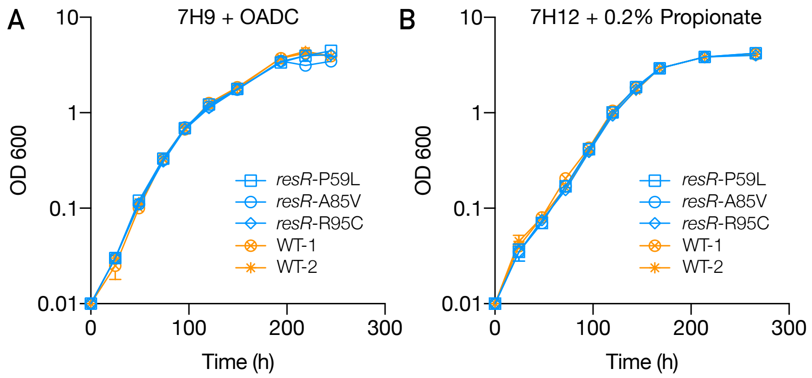


**Fig. S3. Growth curves of *resR* mutants and wild-type *Mtb* strains.** Growth curves of *resR* mutants and wild-type *Mtb* strains under the standard (7H9 + OADC) or a host-relevant (7H12 + 0.2% Propionate) condition. *y* axis is shown in $\log_{10}$ format, and the data represent the mean and standard deviation of two biological replicates and two technical replicates.

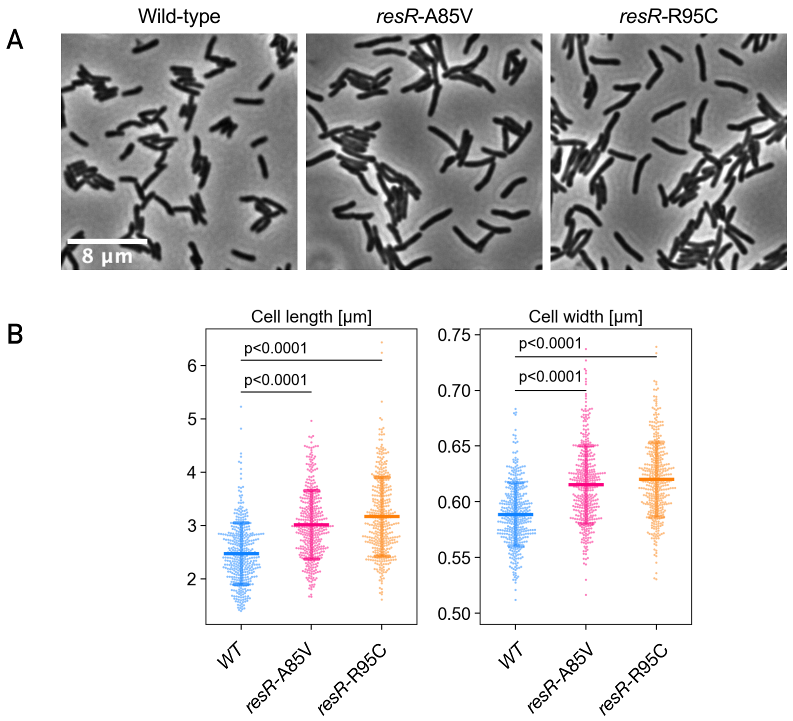


**Fig. S4. *resR* mutants exhibited increased cell lengths and widths.** (A) Representative microscopy images of wild type and two resR mutants. (B) Single-cell length and width profiles of wild type and two resR mutants. Image segmentation and morphological profiling was performed as described in Methods. 400 cells were randomly sampled from each strain's dataset to render a uniform representation of their morphological profiles.

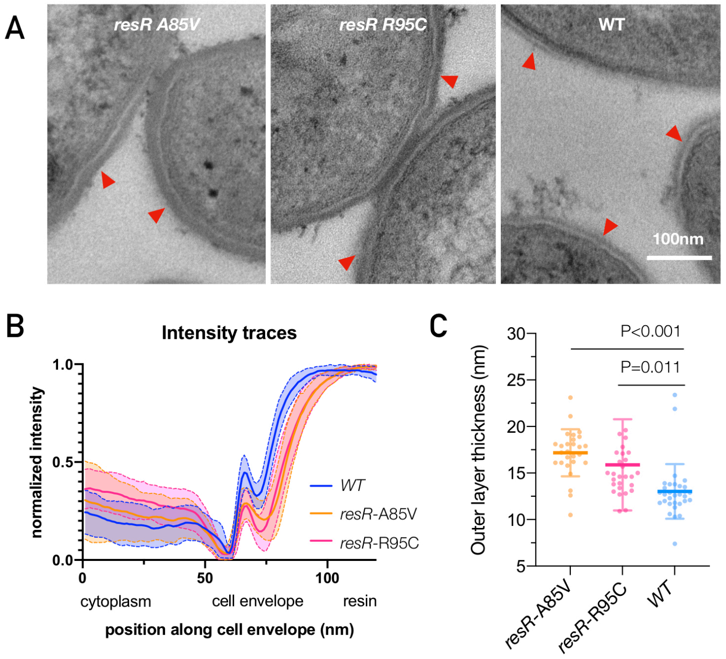


**Fig. S5. Cell envelop intensity traces of *resR* mutants and WT via TEM.** (**A**) Transmission electron microscopy (TEM) images of *resR* mutants and wild-types showing examples of the difference in the thickness of the outer cell envelope. (**B**) Intensity traces used to measure outer cell envelope thickness. Line profiles were drawn perpendicular to the envelope. The intensity was measured along each line, and min/max normalized; these traces were then averaged to create an average intensity profile (solid lines, with standard deviation indicated by shaded regions). (**C**) Comparison of cell wall outer layer thickness (measured via TEM) between *Rv1830* mutants and wild-type strains. 31.7% and 21.9% increase for A85V and R95C respectively. Each dot represents one Mtb cell and P values were obtained by unpaired t test with Welch's. Error bars show standard deviation.

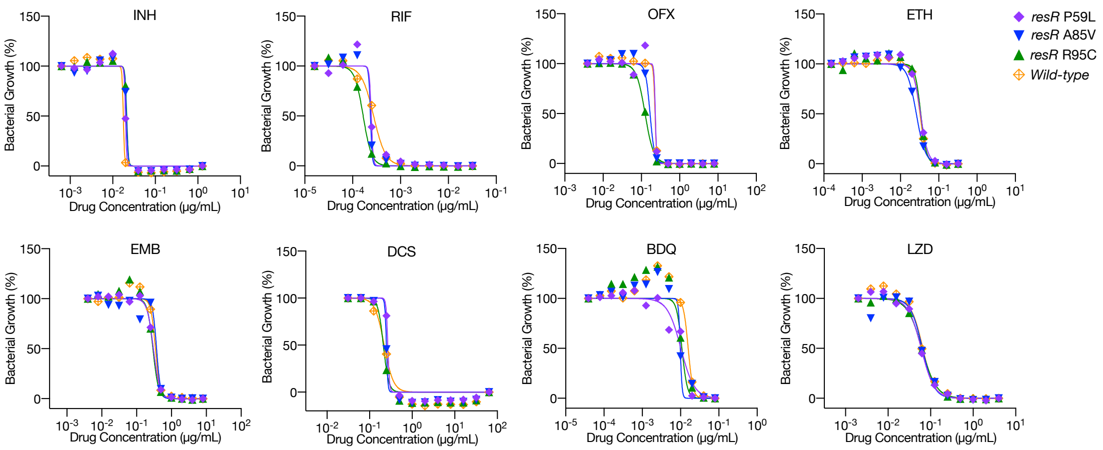


**Fig. S6. Antibiotic susceptibility (MIC) of *Mtb* resR mutants and wild-type strains to 8 anti-tuberculosis drugs**. The experiment was performed three times with similar results.

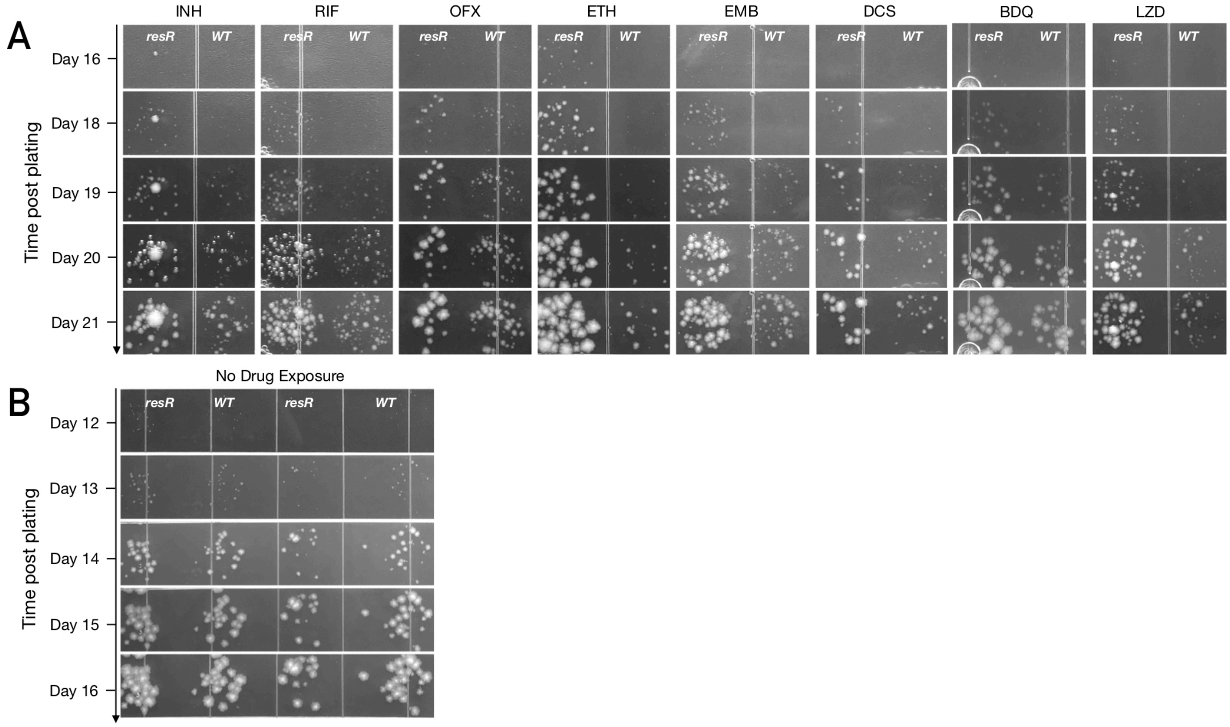


**Fig. S7. *resR* mutants showed earlier appearance of visible colonies post antibiotic exposure.** Representative photo clips illustrating the post-antibiotic recovery dynamics of *resR* mutant and wild type (WT) *Mtb* strains from “Drug exposure” (**A**) and “No drug exposure” (**B**) groups. *Mtb* cells were exposed to 100-fold MIC concentrations of antibiotics for 24 hours and then were washed and seeded onto drug-free agar plates. “Time post plating” denotes the time interval (in days) between plating and photographing.

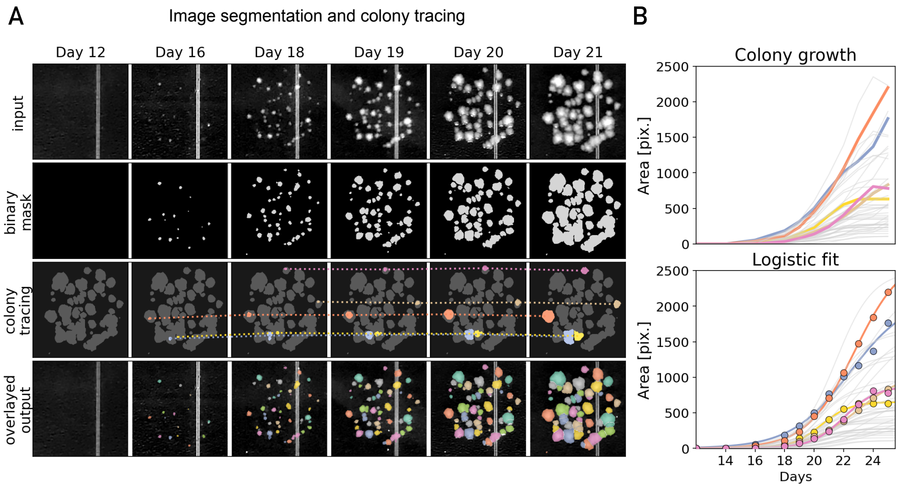


**Fig. S8**. **Schematic overview of automated colony tracing and quantitative image analysis.** (**A**) Pre-processed plate photos were clipped into individual clusters of colonies (top panels), which were then converted into binary masks (mid-top panels) using a pre-trained pixel classifier (Methods). The masked regions were further segmented and linked using a forward-matching algorithm to approximate the growth dynamics of individual colonies. Here 5 representative colonies were pseudocolored, and a dashed line was drawn to link the corresponding masked pixels of different time points for each colony (mid-bottom panels). The final tracking records were pseudocolored and depicted in the bottom panels. (**B**) Line plots representing the raw growth dynamics (top panel) of all segmented colonies in (A) or their logistic-fitted curves (bottom panel) in pixel area units. The curves of the 5 representative colonies depicted in (A) were highlighted and colored correspondingly.

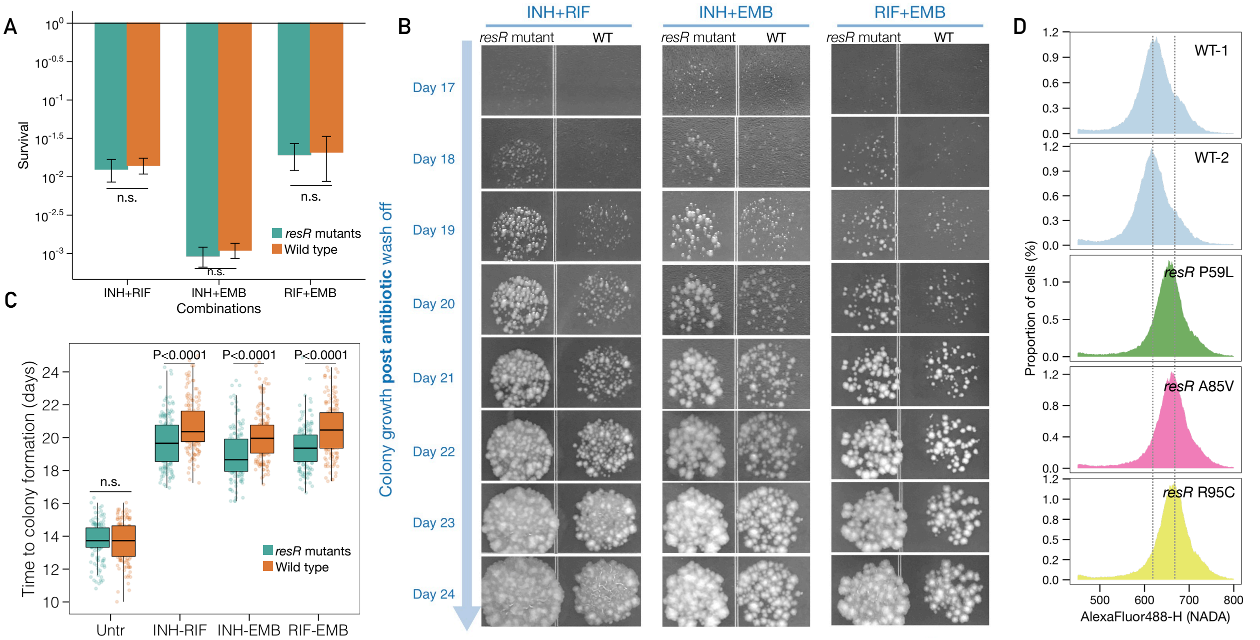


**Fig. S9. *resR* mutants showed faster recovery after treatment by drug combinations.** (**A**) *resR* mutants showed no difference in survival after 24h treatment by different combinations of first-line anti-tuberculosis drugs. The concentration of each individual drug was used as 10-fold of the MIC. (**B**) Aligned photo clips representing the colony growth dynamics of *resR* mutants and wild-type strains after 24h treatment by different combinations of antibiotics. Plate photos were taken every 24 hours. (**C**) A box plot showing the time duration from plating to the appearance of visible colonies (Time to colony formation) for *resR* mutant and wild-type strains after exposure to three different combinations or no drug exposure (Untr). (**D**) Flow cytometry for NADA incorporation into Mtb cells after 24 hours recovery post 100X INH exposure for wild-type and *resR* mutant strains.

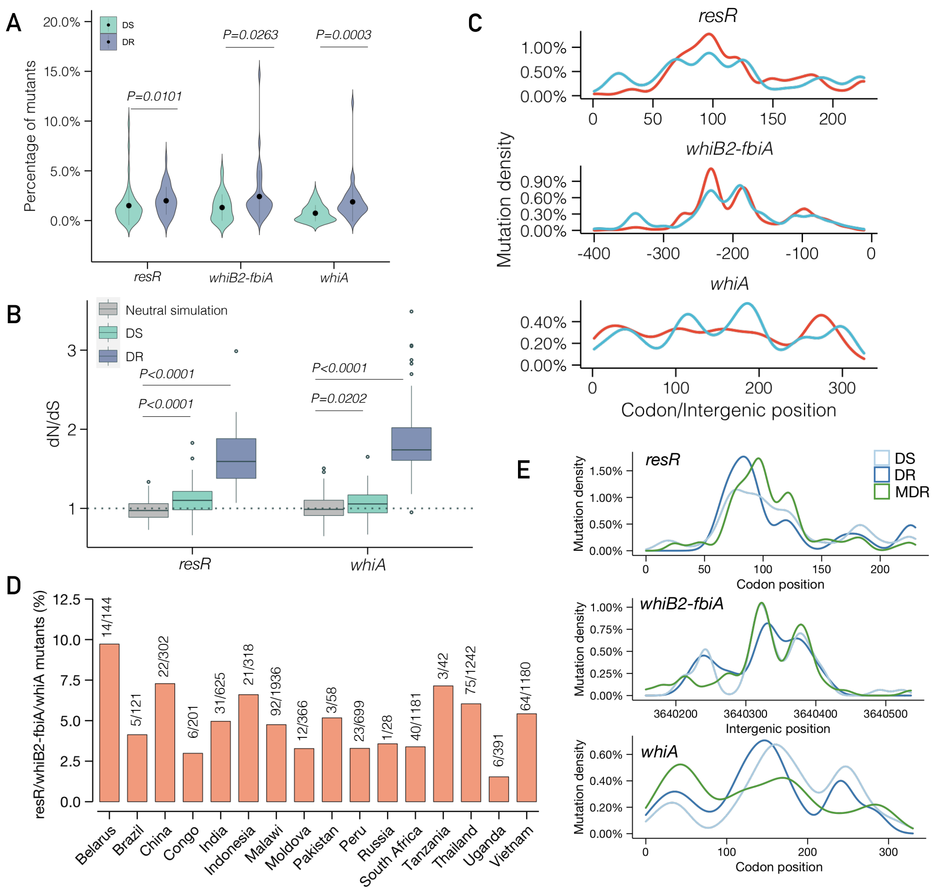


**Fig. S10. *resR, whiB2-fbiA and whiA* mutations are under selection in both DR and DS *Mtb* strains.** (**A**) Percentage of *Mtb* isolates with nonsynonymous mutations in genes of R*v1830,* and *whiA,* and mutations in *whiB2-fbiA* IGR region from all countries' data. (**B**) dN/dS ratio of *Rv1830* and *whiA* in DS, DR and neutral simulation groups. dN/dS ratio was calculated by subsampling 500 *Mtb* isolates from each group and repeated 100 times. Neutral simulation represent the relative dN/dS ratio from a random substitution model. (**C**) Mutational pattern and hotspots of fixed mutations in DR (red) and DS (blue) strains. (**D**) Percentage of DS strains with *resR, whiB2-fbiA* or *whiA* mutations in 16 high TB burden countries. (**E**) Mutational pattern and hotspots of unfixed mutations in DS, DR and MDR-TB strains.
