## Supplementary material for "Tuberculosis treatment failure associated with evolution of antibiotic resilience": Materials and Methods

### *Genome sequences of Mtb isolates*

We used the keywords "tuberculosis", "*Mycobacterium tuberculosis*" and "*Mycobacterium tuberculosis* complex" to search the NCBI Sequence Read Archive (SRA) database for whole-genome sequencing records of *Mtb* clinical isolates. Whole-genome sequencing data for 51,229 *Mtb* isolates from 203 SRA projects were included in this study. The tool *prefetch* (https://www.ncbi.nlm.nih.gov/books/NBK242621/) was used for downloading the *sra* files of the *Mtb* isolates. The geographic information of the *Mtb* isolates was obtained from the BioProjects, published articles or shared by the corresponding authors. In total, geographic information for 46,104 *Mtb* isolates was identified. The downloaded *sra* files were converted to paired-end or single-end *fastq* files using the toolkit of *fastq-dump* (https://www.ncbi.nlm.nih.gov/books/NBK158900/). Samples with an average sequencing depth over 20X and a mapping rate of over 90% were used for downstream analyses.

### *SNPs calling*

Sequencing reads were trimmed with *Sickle (1)*. Trimmed reads with length > 30 and Phred scores > 20 were retained for subsequent analyses. The inferred ancestral genome of the most recent common ancestor of the *MTBC* was used as the reference template for reads mapping (*2*). Sequencing reads were mapped to the reference genome using *Bowtie* 2 (v2.2.9) (*3*). *SAMtools* (version 1.3.1) was used for SNP calling with the minimal mapping quality set to be 30 (*4*). We excluded SNPs located in repetitive regions of the genome (e.g., PPE/PE-PGRS family genes, pro-phage genes, insertion or mobile genetic elements) that are difficult to characterize with short-read sequencing technologies (*5*).

*Unfixed SNPs.* Unfixed SNPs with a frequency of 1%~95% and at least 3 reads supporting the mutant alleles (at least one read from the forward strand and one read from the reverse strand) were identified using *VarScan* (v2.3.9) with the strand bias filter on (*6*). A previously validated pipeline was used to filter out false positives (*7, 8*). First, *mpileup* files were generated by *the samtools-mpileup* function and were used to identify the locations of mutant alleles in the relative sequencing reads, and unfixed SNPs with more than 50% of supporting alleles enriched in the terminal region of sequencing reads were excluded. Second, the sequencing depth of the site with unfixed SNPs should be within 50%~150% of the isolate's average sequencing depth. This is to remove false positives due to INDELs or duplications. Third, all unfixed SNPs that were mapped to repetitive regions of the genome were excluded. Finally, unfixed SNPs with repetitive occurrence but similar mutational frequencies (suggestive of multi-locus mapping) were also removed.

*Fixed SNPs calling.* Fixed SNPs with a frequency of ≥95% and at least 10 supporting reads were identified using *VarScan* (v2.3.9) with the strand bias filter on. Small insertions or deletions (INDELs) identified by *VarScan* (v2.3.9) were also excluded. *Mtb* isolates were typed into lineages based on the previously defined lineage-specific barcode SNPs (*9, 10*). We excluded isolates showing heterozygous typing results or those that were missing lineage-defining SNPs in the lineage-associated analysis.

***Genotypic drug susceptibility test***

As DNA sequencing can accurately predict profiles of antibiotic susceptibility with sensitivity between 91.3%-97.5% for the first-line anti-tuberculosis drugs (isoniazid, rifampin, ethambutol and pyrazinamide) (*11*), we used the previously validated DR mutations for genotypic drug susceptibility testing for *Mtb* isolates. The drug resistance-associated mutations in *Mtb* were obtained from the database TBDReaMDB and articles reporting validated DR mutations (*11, 12*). *Mtb* isolates carrying any previously characterized DR mutation (fixed SNP) were identified as DR strains, whereas the ones with mutations indicating resistance to both isoniazid and rifampicin were determined as MDR-TB strains. The remaining isolates with no known DR mutations were identified as DS strains.

***Mutation burden test***

The unfixed SNPs in the population of a clinical *Mtb* isolate can represent the mutations that accumulated within-host during the infection/treatment, or from mixed infections of multiple *Mtb* strains. Mixed infections of two or more genetically distinct strains would result in large numbers of unfixed SNPs, typically presented as two or more groups of unfixed SNPs with similar SNP frequencies(*13, 14*). We applied a previously developed pipeline to filter suspected samples of mixed infections by excluding isolates with a cohort of unfixed SNPs that exclusively mapped to known evolutionary paths(*14*). Then, we grouped all unfixed mutations from isolates of monoclonal infections and each unfixed SNP was considered as an independent mutation event since they arose in *Mtb* populations isolated from different patients. In total, 240,064 unfixed SNPs were included in the subsequent analyses. *Mutation* *burden test* was conducted by calculating the normalized mutation density of unfixed SNPs on each *Mtb* gene as follows:

$$Di = Mi/(N \times Li)$$

In this equation, *Di* refers to the mutation density of gene *i*, *Mi* refers to the total unfixed SNPs observed for gene *i*, *N* is the total number of unfixed SNPs while $L_i$ is the length of gene *i.*

***dN/dS***

As the ratio of nonsynonymous (*dN*) to synonymous (*dS*) nucleotide substitution is an indicator of selective pressures on genes, we calculated *dN/dS* for each *Mtb* gene to evaluate the selective pressure. A ratio greater than one indicates positive selective pressure (selection favors changes) while a ratio under one indicates purifying selection (pressures to conserve protein sequence)(*15*). All *Mtb* isolates were grouped as a population and the *dN/dS* was calculated to interpret selective pressure on each gene at the population level. For unfixed mutations, the mutation events of each gene were concatenated into one sequence and we used the *kaks* function in R package *seqinr* to calculate *dN/dS(16)*. As the estimation of *dN/dS* for fixed mutations could be affected by recent clonal expansion events or the convergent evolution of a particular site in an *Mtb* population (e.g., DR mutations), we conducted a bootstrap analysis by sub-sampling 1000 isolates from the 51,229 *Mtb* isolates and estimated the *dN/dS* of fixed mutations of the sampled subset. This process was repeated 100 times. To estimate *dN/dS* of the DS, DR and MDR groups, isolates were first assigned to different groups based on their DR status and the *dN/dS* of each group was calculated separately.

To test whether the observed *dN/dS* ratios of *resR* and *whiA* in DS *Mtb* strains would be a significant deviation from neutral evolution (fig. S10B), we used a previously described method to generate mutations *in silico* through a random substitution process (*7*). Briefly, a codon substitution matrix was generated using a base substitution model which considers the genome's GC content (65.6% for *Mtb*) and the proportion of transitions that occurred at the wobble position of codons in synonymous fixed mutations (Ti, 0.729). For each codon, a simulation of 50,000 individual introductions of a single mutation was performed and the outcomes were scored as either synonymous or nonsynonymous. The average number of nonsynonymous outcomes of the simulations is an estimate of the probability that a mutation in the given codon would be nonsynonymous.

***Protein structure modeling***

The model of the ResR (Rv1830) protein structure was obtained from AlphaFold (https://alphafold.ebi.ac.uk). The AlphaFold Model Confidence value (pLDDT) for the MerR-type helix-turn-helix (HTH) DNA-binding domain of ResR is >90 for nearly all residues. Using PyMOL (Schrodinger), the model of ResR was aligned to the crystal structure of the activator form of *E. coli* CueR (a merR-type regulatory protein homolog) in complex with a 23 base pair strand of duplex DNA based on the *E. coli copA* promoter (PDB: 4WLW) (*17*). The alignment has a Root Mean Square Deviation (RMSD) of 1.019. One copy of ResR was aligned to each copy of the CueR dimer to generate the dimeric ResR DNA-bound model.

***resR point mutants construction***

*resR* mutants were constructed using oligo-mediated recombineering as previously described (*18*). First, the *Mtb* H37Rv strain was transformed with two plasmids, pKM427 and pKM402. pKM427 is a chromosomal site L5 integrating vector carrying a zeocin resistance cassette and a hygromycin cassette that is inactivated by a premature stop codon, while pKM402 is an episomal vector carrying the phage *recT* recombinase gene under an ATc inducible promoter. The resultant *Mtb* strain was grown to early log phase, induced with ATc to express RecT, then transformed with two single-strand DNA oligos, with one oligo conferring hygromycin resistance and the other oligo carrying one of the three clinical *resR* mutations (P59L, A85V and R95C). The transformants were recovered for 2 days in drug-free media (7H9 with 0.2% glycerol, 10%OADC supplement, 0.05% Tween80) and plated on solid media (7H10 agar, 0.2% glycerol, 10% OADC supplement, 0.05% Tween80) containing hygromycin (50 µg/ml). Colonies were screened for *resR* mutations by Sanger sequencing and two colonies were retained for each mutant of interest. In parallel, we obtained three hygromycin-resistant colonies from separate transformations with no *resR* mutations and used them as the wild-type controls. All strains were expanded and re-streaked onto solid media with hygromycin and colonies which had spontaneously cured the pKM402 plasmid were saved and used for downstream experiments.

The *Msmeg* T77A (*MSMEG_3644*) mutant strain was constructed using oligo-mediated recombineering in *Msmeg* mc$^2$155 strain as described above, but the chromosomal site L5 integrating vector contained a kanamycin resistance cassette and transformed cells recovered for 3 hours in drug-free media. The *Msmeg MSMEG_3644* CRISPRi knocking-down strain was constructed as previously described (*19*). The forward *MSMEG_3644* sgRNA targeting oligo was 5’-3’ GGGAGCCGCCATCGGCGCGCTCGCT and the reverse oligo was

AAACAGCGAGCGCGCCGATGGCGGC, which correspond to the PAM sequence CGGGAAG. The complimentary oligos were ligated into BsmBI-digested PLJR962, the Sth1 dCas9 CRISPRi vector backbone. This construct was transformed into *Msmeg* $mc^2155$ strain and transformants were recovered after plating on solid media with kanamycin (20 µg/ml). We confirmed MSMEG_3644 knockdown via qPCR after overnight induction with ATc (100 ng/mL) in media with kanamycin (20 µg/ml). The *Msmeg MSMEG_3644* overexpression strains were constructed by transforming the *Msmeg* $mc^2155$ strain with a vector containing wild-type or *MSMEG_3644* T77A under the control of a Tet repressor-regulated UV15-Tet promoter, a kanamycin selection marker, and a single-copy L5-integration site (the T77A mutation was introduced into the vector expressing wild-type *MSMEG_3644* via PCR site-directed mutagenesis. The transformants were recovered after plating on solid media with kanamycin (20 µg/ml). Overexpression of wild-type and *MSMEG_3644* T77A was verified with qPCR after overnight induction with ATc (100 ng/mL) in media with kanamycin (20 µg/ml).

***Transmission Electron Microscopy***

*Mtb* cultures of *resR* mutants and wild-type strains were grown to late log phase ($OD_{600}$ around 1.0) and the cells were pelleted via centrifugation at 4,000xg at room temperature for 10 min, then resuspended in fresh 7H9 media. The suspension was subsequently combined with an equal volume of 0.1M sodium cacodylate buffer (pH 7.4) containing 1.25% formaldehyde, 2.5% glutaraldehyde, and 0.03% picric acid. The cells were fixed at room temperature for 2 hours, then washed with sodium cacodylate buffer three times to remove the fixatives. The cell pellet was stained in 1% osmium tetroxide for 1 hour at room temperature, washed with deionized water three times, then stained with 1% uranyl acetate for 1 hour. The stained samples were washed three times with deionized water then dehydrated by washing with 70% ethanol once, 90% ethanol once, and 100% ethanol twice. The ethanol-dehydrated samples were further treated in propylene oxide for 1 hour at room temperature, then immediately subjected to resin infiltration by immersing in a 1:1 mixture of Spurr'sresin and propylene oxide overnight at 4 ˚C. After infiltration, the samples were transferred to the embedding mold filled with freshly mixed Epon and allowed to polymerize at 60 ˚C for 24 hours. Ultrathin sections (about 80nm) were cut on a Reichert Ultracut-S microtome, picked up onto copper grids stained with lead citrate, and examined in a JEOL 1200EX Transmission electron microscope images were recorded with an AMT 2k CCD camera.

Outer cell envelope thickness measurements were performed using a custom-built MATLAB (Mathworks) script. All images were downsampled to 1 nm/pix. Image intensity profiles were extracted perpendicular to a user-input drawn line along the cell membrane. The distance between the middle cell envelope layer (defined as the highest peak between 55 nm and 75 nm from the beginning of the line profile) and the inflection point of the curve between the outer cell envelope layer and the resin (defined as the highest point in the image gradient between 70 nm and 100 nm from the beginning of the line profile) was measured as the outer cell envelope thickness. This experiment was conducted once, and the average outer cell envelope thickness for each cell was recorded. The line intensity profiles were min/max normalized and averaged to produce an average electron density profile for each strain measured.

***MIC experiments***

Minimum inhibitory concentrations (MICs) for 8 anti-tuberculosis drugs (INH, RIF, OFX, ETH, EMB, DCS, BDQ and LZD) were measured using the Alamar Blue reduction assay. Briefly, strains were grown to mid-log phase ($OD_{600}$ ~ 0.5) and diluted to a final $OD_{600}$ of 0.003 in 7H9 media (with 0.2% glycerol, 10%OADC supplement, 0.05% Tween80) containing antibiotics at indicated concentrations. After five days of incubation at 37˚C with constant agitation (60 rpm), Alamar Blue reagent was added to each well and incubated at 37˚C for two more days before measuring the cultures' absorbance profiles. The amount of Alamar Blue reduction (denoted reduction amplitude or RA) was estimated by subtracting background OD at 600nm from OD at 570nm. The percentage of bacterial growth was calculated by firstly subtracting the RA measure of the no-cell well (baseline) and then scaling the residue by RA of the no-drug control (maximal growth). The growth inhibition dynamics were approximated by linear regression on the log2-transformed fold-change data using Prism 9. The minimum inhibitory concentration of each drug was determined based on two independent experiments with each experiment containing two replicate sets of wells.

***Time-kill assays***

*Mtb resR* mutant and WT strains were cultured in 7H9 media (with 0.2% glycerol, 10% OADC supplement, 0.05% Tween80) to mid-log phase ($OD_{600}$ ~ 0.5) and then diluted to a final $OD_{600}$ of 0.05. Two parallel cultures were prepared for each strain and 10 mL of each diluted culture was inoculated into a 50 mL inkwell. Antibiotics were added to final concentrations of 100 × MIC of the *Mtb* strains (INH: 1μg/mL for WT, 2 μg/mL for *resR* mutant; RIF: 0.2 μg/mL; OFX: 25 μg/mL; ETH: 10 μg/mL; EMB: 50 μg/mL; DCS: 400 μg/mL; BDQ: 2 μg/mL; LZD: 25 μg/mL). The antibiotic-supplemented cultures were incubated at 37˚C with constant agitation (60 rpm), and sampled at day 1, day 3, and day 5 to plate for colony enumeration. Briefly, 1 mL aliquot of each antibiotic-supplemented culture was centrifuged at 6000 rpm for 5 min at room temperature, then the supernatant was removed, and the pellet was resuspended in 1 mL fresh 7H9 media (with 10% OADC supplement, 0.2% Glycerol, 0.05% Tween80) with no antibiotics. An additional round of centrifugation-wash cycle was performed to remove the residual antibiotics in the media. After washing off the antibiotics, a series of 5-fold dilutions of the washed cultures were prepared in a 96-well plate using fresh 7H9 media, and 4 μL of each dilution was spotted on a 7H10 agar plate with no antibiotics. Colony enumeration was performed at 24 days after plating, and the dilutions that yielded 5~50 visually separated colonies were counted and used for survival estimation.

***Quantitative imaging of colony growth***

To track colony growth dynamics, antibiotics-challenged or untreated *Mtb* cultures were washed according to the procedures described above and spotted onto plain 7H10 agar. Plate photos were taken using an 8-megapixel camera at 12, 13, 14, 15, 16, 18, 19, 20 days after plating for cultures without antibiotic challenge; or 12, 13, 14, 15, 16, 18, 19, 20, 21, 22 days for cultures treated with a single drug; or 17, 18, 19, 20, 21, 22, 23, 24, 25, 26, 27 days for cultures treated with a combination of first-line anti-tuberculosis drugs. We devised an image processing pipeline to realign and transform the plate images into a uniform shape aligning by the grids of the plates. Briefly, the plate photos were first converted into 8-bit grayscale images using Fiji software, then processed with a low-frequency bandpass filter to suppress background illumination variations. The coordinates of 49 key points (intersection points of the 7×7 grid lines of the square petri dish) of each bandpass filtered image were manually determined using Fiji's ROI tools, these key points were then used to guide the projection of the distorted photo onto a standard image space of 2000 × 2000 pixels using a piece-wise affine transform algorithm. For each plate, we estimated the relative lateral drifts over time by their cross-correlation in frequency space and realigned the time sequence accordingly. Rectangular regions of interest (ROIs) were defined manually using Fiji and used to crop the large plate image sequence into colony clusters, with each cluster corresponding to a specific dilution of a given sample.

To separate pixels of *Mtb* colonies from its complex plate background, we devised a Random-Forrest-type pixel classification model using image textural and spatial features of multiple scales as inputs. The model was trained on a manually annotated dataset of 32 plate image clips with an average of ~20 colonies per clip and achieved over 96% pixel classification accuracy when tested on an independent validation dataset. Using this pre-trained model, we converted the cropped time-series images into binary masks, which were subsequently segmented into interconnected pixel patches and labeled with unique identifiers.

To track the expansion dynamics of a single colony over time, we took a forward matching approach. In brief, a pixel patch was accepted as a colony or the union of multiple colonies if its corresponding pixels overlapped with at least one patch found in the next time frame. In cases where residual drifts exceeded colony sizes, we calculated the geometric centers (centroids) of pixel patches of the two consecutive frames, and iteratively determined the planar offsets that minimize the average distance between the centroids of a given pixel patch and its nearest neighbor found in the next time frame. Patch pairs with an overlap ratio over 0.4, or an offset-corrected centroid distance less than 1 pixel were subsequently linked. Patches that are linked to 2 or more patches of the previous time frame were further split using the watershed method and with its linked patches as the initial seeds. The forward matching – linking – splitting process was repeated until all patches were linked to only one other patch of the previous or the following time frame. A uniquely linked patch series therefore corresponded to a single colony, and the areas (pixel unit) of these patches were used to approximate this colony's growth dynamics.

***Modeling of colony growth dynamics***

Colony expansion rate is defined as:

$$f_{expansion}(t, dt) = \frac{A_t - A_{t-dt}}{dt \times A_{t-dt}}$$

Here $A_t$ denotes the colony's area measured at $t$ days after plating, whereas $dt$ denotes the interval between two consecutive sampling times. The maximum expansion rate of a given colony is defined as the numeric average of the three highest expansion rate measures. Colonies with a maximum expansion rate over 2.5 is considered abnormal and excluded for downstream analysis.

To estimate the time to colony formation, we first modeled the colony's growth dynamics using a generic logistic function:

$$g(t) = \frac{A_{max}}{1+e^{-k(t-t_0)}}, 0 \leq \ t \leq 30(days)$$

Here the three parameters $A_{max}$, $k$, and $t_0$ denote the maximum colony area, the logistic growth rate and the time required to reach half-maximum area. For each colony, we fitted the above function to its area measures over time using a non-linear least squares method. The estimated time to colony formation was therefore defined as the value of $t$ when $g(t)$ surpasses an arbitrarily defined threshold of 50 pixels.

***Pulse-chase experiment***

Dual-color fluorescent D-amino acid (FDAA) pulse-chase labeling was conducted as previously described by Botella et al.(*20*). Late log phase ($OD_{600}$ ~1) cultures of WT or *resR* mutant strains were sub-cultured in fresh 7H9 media (with 0.2% glycerol, 10%OADC supplement, 0.05% Tween80) supplemented with 25 μM NADA (3-[(7-Nitro-2,1,3-benzoxadiazol-4-yl) amino]-D-alanine hydrochloride) at a final $OD_{600}$ of 0.05. The NADA-containing suspensions were aliquoted in a non-tissue culture treated 24-well plate (1 mL culture each well) and incubated at 37˚C for two days with constant agitation until the cultures had reached $OD_{600}$ of 0.2~0.3 and the cells had fully incorporated NADA into their peptidoglycans. INH was added to the antibiotic exposure groups to a final concentration of 1μg/mL (100 × MIC). After 24 hours of antibiotic treatment, *Mtb* cells of each well were pelleted by centrifuging at 6000 rpm for 5 min at room temperature, and the pellets were resuspended in fresh 7H9 media. The washing step was repeated to remove the residual INH or NADA molecules. The washed cultures were then inoculated into new 24-well plates containing 7H9 media and 25 μM of a second FDAA, HADA (3-[[(7-Hydroxy-2-oxo-2H-1-benzopyran-3-yl) carbonyl] amino]-D-alanine hydrocholoride), and incubated at 37˚C with constant agitation. At 10, 24, or 36 hours after re-inoculation, one 24-well plate replicate was harvested and the cultures were mixed with an equal volume of 4% PFA and incubated at room temperature for 2 hours. The PFA-fixed *Mtb* cells were transferred out of the BSL-3 biosafety facility and stored at 4˚C for no more than 72 hours before microscopy.

***Flow cytometry***

To rapidly quantitate the post-antibiotic growth recovery of *resR* mutant and WT strains via flow cytometry, we adopted a similar bacterial culture procedure as the pulse-chase experiment except that no NADA was added before the antibiotics challenge. After the cultures had reached $OD_{600}$ of 0.2~0.3, antibiotics were added to the culture at 100 × MIC equivalent concentrations. After 24 hours of antibiotics challenge, cells were washed twice with an equal volume of plain 7H9, then resuspended in 7H9 supplemented with 25 μM NADA. Post-antibiotics NADA incorporation was carried out for another 24 hours before the cells were fixed and removed from the BSL-3 biosafety facility as described above. Fixed bacilli were quenched with 200mM Tris-HCl (pH 7.5) for 5 minutes at room temperature and resuspended in PBST buffer (1×PBS supplemented with 0.1% TritonX-100). Quantitation of NADA incorporation was conducted on a MACSQuant® Analyzer 10 flow cytometer using the green fluorescence laser and filter set (channel B1, Ex. 488nm and Em. 525/50nm). As *Mtb* cells are small and prone to cellular aggregation due to their hydrophobic cell surfaces, both pre- and post-acquisition quality control measures were taken to suppress faulty events. To suppress signals from noise or cell debris, two event triggers (thresholds) on forward scatter peak height (FSC-H >1.5) and side scatter area (SSC-A > 1.0) were used upon recording. To remove cellular aggregation, stringent gate settings were manually defined via FlowJo v. 10.8 to exclude events with strongly correlated forward scatter area (FSC-A) and SSC-A measures (large and compact particles), as well as events with disproportional FSC-A and FSC-H measures (morphological outliers). After event filtration, the log10-transformed green fluorescence intensity peak height (denoted AlexaFluor488-H) was used to represent the post-antibiotic cell wall synthesis activity of *Mtb* strains.

***Quantitative microscopy***

For each microscopy run, 100 µL PFA-immersed *Mtb* suspensions were centrifuged at 3,000 × g for 15 minutes, the pellets of which were dissolved in 200 µL of customized cocktail containing 200mM Tris-HCl (pH = 7.5), 1% (w/v) Triton X-100, 0.67% (v/v) Xylenes and 0.33% (v/v) Heptane. This step was found to efficiently reduce cellular aggregation and quench the remaining aldehydes. The disaggregation-quenching procedure was carried out for no more than 2 minutes, after which the cells were pelleted by centrifugation at 3,000 × g for 15 minutes, washed with 200 µL PBS-Tx (PBS containing 0.1% Triton X-100), then resuspended in 20 µL PBS-Tx. 0.5 µL aliquots of the reconstituted cell suspensions were spotted onto a custom-made 96 well agarose pedestal arrays (1.8% agarose in 1X PBS, supplemented with 0.1 µL/mL Nile Red). The assembled imaging cartridge was incubated at 37C for 15 minutes for Nile Red staining to reach equilibrium. Images were acquired using a Nikon Ti-E inverted, widefield microscope equipped with a Plan Apo 100× 1.45 NA objective lens, a Nikon Perfect Focus system with a Piezo Z drive motor, an Andor Zyla sCMOS camera, and the NIS Elements software (v4.5). Automated high-throughput imaging was carried out using a customized Nikon JOBS script to locate imaging fields of interest, a total of 24 images were taken for each sample. Fluorescent signals were acquired using a 6-channel Spectra X LED light source and the Sedat Quad filter set. The excitation (Ex.) and emission (Em.) filters used in this study were: Ex. 395/25nm and Em. 435/25nm for HADA; Ex. 470/24nm and Em. 515/25nm for NADA; Ex. 550/15nm and Em. 595/25nm for Nilre Red. An exposure time of 100ms and a laser power at

100% were used for all three channels. Bacterial segmentation and morphological profiling was conducted using our previously established microscopy image analysis pipeline, MOMIA (ref). Segmented cells were further filtered using a particle classification model pretrained on >70,000 manually labeled instances to remove incorrectly segmented particles. Single-cell axial fluorescence profiles were measured along the cellular centerline and were used to estimate cell pole growth after antibiotic treatment. A bacterium's polar outgrowth is defined as the length of terminal segments with NADA signals lower than 45% of the cell's maximum NADA staining signal.

***ResR protein purification***

The DNA sequence of wild-type *resR* was cloned into the expression vector pET28b with either a N-terminal 6xHIS tag or a C-terminal 6xHIS tag, the resultant plasmids were transformed into BL21 CodonPlus (DE3)-RP cells (Agilent, catalog 230255) separately. The BL21 strains harboring ResR expression plasmid were cultured in 200 mL of LB media with 50 ug/ml kanamycin with shaking at 28°C. At $OD_{600}$ = 0.1, protein production was induced with 0.1 mM IPTG overnight. Cells were pelleted at 5000 x g and resuspended in HIS-Pur buffer (50 mM Tris-HCl, pH 8.0, 100 mM NaCl, 50 mM KCl) before freezing. Frozen pellets were thawed and resuspended in 10 mL lysis buffer (HIS-Pur buffer with protease inhibitor and lysozyme), then lysed via sonication for 3 x 30 seconds with 30% amplitude and 50% duty cycle (equipment cat#). The samples were kept on ice during sonication to avoid overheating. The lysates were transferred to a cold room and all subsequent steps were performed at 4°C. The lysates were spun at 12,000 x g for 20 minutes and the supernatants were combined with 1 mL of Ni-NTA Agarose (QIAGEN) and incubated overnight with slow shaking. Then, samples were loaded to a gravity flow column and allowed to drain. The samples were washed with 1 mL wash buffer (HIS-Pur buffer with 20 mM imidazole) and eluted with elution buffer (HIS-Pur buffer with 100 mM imidazole). Elution fractions were analyzed by SDS-page and protein concentrations were measured by Qubit 3.0. Eluted ResR proteins were dialyzed in 1 mL dialysis buffer (0.05mM Tris pH 8.0, 0.3M NaCl, 0.1% Glycerol, 1mM DTT, 0.5mM EDTA) using 1 mL dialysis cassettes (Thermo Scientific) at 4°C overnight, and then stored at -80°C.

***In vitro DNA-protein binding sequencing (IDAP-seq)***

*Mtb* genome DNA was sheared to an average fragment size of 300bp (Covaris E220e at 140 W, a duty factor of 10%, 200 cycles per burst, and an 80 second treatment time). ResR proteins were thawed on ice 2 hours before being combined with 3.4 μg sheared DNA and 4 μM ResR protein in a final volume of 125ul with binding buffer (20 mM Tris pH 8.0, 300 mM NaCl, 0.2 mg/mL bovine serum albumin, 10% glycerol, 1 mM dithiothreitol, and 200 ng of sheared herring sperm DNA). Binding reactions were incubated for 30 minutes at room temperature, and then mixed with a 50 µL bed volume of Talon resin (Clontech) pre-equilibrated in binding buffer (20mM Tris pH8.0, 300 mM NaCl, 010% glycerol, 1 mM dithiothreitol) and incubated for an additional 30 minutes at room temperature with rotation. Bound ResR-resin mixtures were then loaded onto Poly-Prep chromatography columns (Bio-Rad). Columns were washed 3 times with 1 mL of binding buffer. To elute ResR-bound genomic DNA, 500 µL of pre-warmed elution buffer (50 mM Tris pH 8, 10 mM EDTA, 1% SDS) was added to the resin and the column was capped and sealed with parafilm to minimize evaporation, then incubated on a heat block pre-heated to 65˚C for 15 minutes. After incubation, elutions containing ResR-bound DNA fragments were collected and were brought to 1 mL with nuclease-free water, then mixed with 100 µL of AMPure XP beads pre-washed and re-suspended in custom precipitation buffer (20% w/v PEG-8000, 2.5 M NaCl). A fraction of the IDAP-seq sample was purified using the standard AMPure XP dsDNA purification kit according to the manufacturer's instructions. Libraries were prepared using the NEBNext Ultra II DNA Library Prep Kit for Illumina sequencing platform following manufacturer guidelines. Libraries were sequenced on a MiSeq instrument (Illumina) to yield 75 nt / 75 nt paired-end reads.

***RNA-Seq***

*Mtb* or *Msmeg* strains were grown in liquid media (7H9 with 0.2% glycerol, 10%OADC supplement, 0.05% Tween80) to mid-log phase ($OD_{600}$ around 0.5). For each strain, 5 mL of culture was spun down at room temperature and the cell pellet was resuspended in 1mL Trizol lysed via bead beating then combined with 300ul of chloroform. *Mtb* samples were removed from BSL-3 facility after the addition of chloroform. Total RNA was isolated using the Quick-RNA Miniprep Kit (Zymo Research) following the manufacturer's instructions. Ribosomal RNA depletion was done using the Kapa RiboErase kit with custom-made *Mtb*/*Msmeg* rRNA targeting oligos. RNA sequencing libraries were prepared using the KAPA RNA HyperPrep Kit according to instructions for 500ng of input mRNA (Kapa Biosystems). RNA libraries were sequenced on an Illumina MiSeq sequencer with a 75bp single-ended setup. Sequencing reads were then mapped to cognate genomes reference and reads mapping to ribosomal RNAs were removed. For *Mtb* samples, rRNA accounted for less than 1% of the total reads.

For *Msmeg* samples, rRNA accounted for less than 30% of the total reads. Transcriptional level (FPKM) from each gene was identified by the TopHat and Cufflinks packages [ref].

***Phylogenetic reconstruction***

For phylogenetic reconstructions, all SNP locations for each isolate were combined into a non-redundant consensus list and recalled with the mpileup2cns function of *VarScan* (version 2.3.9) [ref]. Nucleotide positions with missing calls in more than 5% of the isolates were removed. An alignment of the remaining polymorphic positions from all strains was used for phylogeny reconstruction with *megacc* [ref]. For the estimation of phylogenies, the Maximum Likelihood (ML) method was applied under the general time reverse (GTR) model with at least 100 replicates for bootstrapping confidence levels. Phylogeny trees were exported from *megacc* and visualized in *FigTree* (version 1.4.3) (http://tree.bio.ed.ac.uk/software/figtree/). We adapted a recently described hierarchical nomenclature to define sublineage nodes and sub-clades within the tree[ref].
